# AgRP neurons coordinate the mitigation of activity-based anorexia

**DOI:** 10.1101/2021.09.10.459764

**Authors:** Ames K. Sutton, Sean C. Duane, Ahmed M. Shamma, Anna Skillings, Michael J. Krashes

## Abstract

Anorexia nervosa (AN) is a debilitating and deadly disease characterized by low body mass index due to diminished food intake, and oftentimes concurrent hyperactivity. A high percentage of AN behavioral and metabolic phenotypes can be replicated in rodents given access to a voluntary running wheel and subject to food restriction, termed activity-based anorexia (ABA). Despite the well-documented body weight loss observed in AN human patients and ABA rodents, much less is understood regarding the neurobiological underpinnings of these maladaptive behaviors. Moreover, while exercise has been shown to diminish the activity of hunger-promoting hypothalamic agouti-related peptide (AgRP) neurons, much less is known regarding their activity and function in the mediation of food intake during ABA. Here, feeding microstructure analysis revealed ABA mice decreased food intake due to increased interpellet interval retrieval and diminished meal number. Longitudinal activity recordings of AgRP neurons in ABA animals revealed a maladaptive inhibitory response to food. We then demonstrated that ABA development or progression can be mitigated by chemogenetic AgRP activation through the reprioritization of food intake (increased meal number) over hyperactivity. These results elucidate a potential neural target for the amelioration of behavioral maladaptations present in AN patients.

## Introduction

Anorexia nervosa (AN) is a global psychiatric disease primarily defined by decreased food intake and low body weight^1^. A key neurobiological feature of AN is the reversal of satiety’s reinforcement value. In healthy individuals, hunger signals a negative valence, which is only relieved upon food consumption. Conversely, AN patients prefer a state of caloric deficiency over sated conditions, and food intake is aversive^2–6^. Current therapeutic strategies for AN patients are primarily behavioral, as no FDA-approved medications exist for the condition, and severe cases oftentimes require forced feeding during prolonged involuntary hospitalization^7–10^. Given the aversive nature of food intake in AN patients, it is perhaps no surprise that relapse rates (~50%) are among the highest of any psychiatric illness; AN is also one of the most lethal psychiatric illnesses, with a standardized mortality ratio around 6^11^. Moreover, AN is more commonly diagnosed in females^12,13^. Thus, there is a huge need for therapeutic alternatives that target the basic biology of AN.

Strikingly, a large proportion (up to 80%) of AN patients also display hyperactivity, due to excessive exercise and/or general restlessness^14–17^. A number of these key features of AN, including many of the sex differences^18^, can be replicated in the activity-based anorexia (ABA) rodent model, in which some animals voluntarily become hyperactive in contexts of food restriction^19–21^. This leads to rapid health deterioration and ultimately death absent experimental intervention. Like humans displaying AN, ABA rodents over-expend energy, potentially due to an excessive drive to forage for food in contexts of deprivation^21,22^. Additional hypotheses suggest that exercise during ABA (also known as starvation-induced hyperactivity) becomes positively reinforcing, potentially to relieve the negative effect of hunger (which later switches to an appetitive drive)^20,23^. Yet, few studies have been able to untangle the neural representation of feeding during ABA, and even fewer have identified neural targets for the amelioration of this disease.

Allelic variability at the level of the *Agrp* gene, the gene encoding the hunger-promoting AGRP peptide in neurons of the arcuate nucleus (ARC), has been identified as a risk factor for AN in humans^36,37^. Surprisingly, plasma AGRP levels are elevated in AN patients, and this is recapitulated at the mRNA and protein level in rodents demonstrating ABA^38–42^. Further complicating matters, chronic intracerebroventricular AGRP infusion ameliorates ABA by increasing feeding and decreasing hyperactivity in rats^41,43^. While AgRP neurons would theoretically provide a unique vantage point for the dissection of these seemingly opposing behavioral adaptations, recent work investigating the role of this population in ABA progression underscores the complexity of the AgRP population during body weight loss.

Historically, AgRP neurons have repeatedly been shown to promote robust food seeking and subsequent food intake across a variety of environmental and physiologic conditions^24–29^. Furthermore, AgRP neurons are inhibited following caloric intake in hungry mice or following bouts of exercise^30–34^. However, activation of AgRP neurons during ABA without food present further promotes hyperactivity in mice without altering future food intake, and thus does not rescue body weight loss^34^. These recent findings call into question the capability of this neural population to ameliorate ABA progression. Yet, it is still unknown whether AgRP activation during contexts of food availability (and thus more similar to the behavioral choices made by AN patients) can alter ABA outcomes. Taken together, it is possible that the elevation of AGRP levels during ABA might be the main driver of hyperactivity in contexts without food present, and that the amelioration of ABA is dependent on specific AgRP activation timing.

We sought to clarify the role of AgRP neurons in ABA progression by first expanding our understanding of the behavioral abnormalities (particularly in feeding behavior) observed in mice displaying ABA. We then tested whether AgRP neurons are capable of both appropriately detecting food intake during food restriction as well as driving feeding behavior to mitigate the excessive body weight loss observed in ABA, with a focus on probing and manipulating AgRP neurons during food availability. To this end, we employed a combination of tools including multi-day, freely-moving *in vivo* fiber photometry and chemogenetic perturbations of AgRP neurons with food available. We discovered that AgRP neurons are functionally dysregulated in response to food intake in ABA conditions, further highlighting this population of neurons as a putative target for ABA maladaptive behavioral choice. Furthermore, artificial activation of AgRP neurons relieves the excessive body weight loss observed during ABA by increasing meal number, leading to increased food intake and reducing physical activity. Importantly, AgRP activation was performed during food presentation, highlighting the capability for AgRP neurons to promote appropriate energy-related choice behavior, a predominant maladaptation in AN patients. Together, these findings suggest a potential therapeutic target for the relief of the hypophagia and hyperactivity observed in AN patients.

## Methods

### Animals

C57BL/6J and AgRP^tm1(cre)Lowl^/J (stock no. 012899) mice were used. Mice were housed with a 12-h light/dark cycle and provided *ad libitum* access to food (standard chow, Envigo 7017 NIH-31, or 20mg grain pellets, TestDiet 5TUM) and water, unless otherwise noted. All mice were group housed until experiments began, after which mice were single housed. All experiments were carried out in adult (>8 weeks) female mice. Some measurements were carried out in the same mouse across conditions, including mice used in chemogenetic (some mice used in both Fig. 4 and Fig. 5), and photometry experiments (e.g. in Activity as well as ABA cohorts). All animal protocols and procedures were approved by the US National Institutes of Health Animal Care and Use Committee.

### Behavioral paradigm

All mice were single housed and separated into one of three behavioral groups (Activity, FR, ABA) with *ad libitum* food and water and access to a voluntary wireless running wheel (Med Associates, ENV-047) in the unlocked (Activity, ABA) or locked (FR) configuration 3-5 days prior to the beginning of experiments. Beginning on Day 0, food was measured and removed from FR and ABA mice 3 hours after the start of the dark cycle. On all days following, food was provided for the first 3 hours of the dark cycle, and food intake was measured. For mice in the Activity cohort, food was measured but not removed. Food provided was standard chow unless otherwise noted. Mouse body weights were measured immediately prior to the start of the dark cycle (prior to food presentation) beginning on Day 0 and throughout the rest of the behavioral paradigm. Running behavior on the voluntary running wheels was initially collected and analyzed using Wheel Manager and Wheel Analysis (Med Associates), respectively.

### Food Intake from Feeding Experimentation Devices (FED3)

In some experiments, detailed feeding information was collected using FED3 devices, which dispense 20mg pellets of chow food either *ad libitum* (Free Feed mode) or at predefined times (Timed Feed mode) ^44^. FED3 devices were made with a 3D Printer (Onyx Pro, Markforged) using a mix of Onyx composite filament and fiberglass (Markforged), pre-made electronics (Open Ephys), and a steel feeding nose (Shapeways). In these experiments, mice were habituated to the FED3 device in the homecage for at least one day in combination with their normal standard chow on the ground of the homecage. After FED3 habituation, chow was removed from the homecage and mice were exclusively fed from FED3 in Free Feed mode. Beginning on Day 0, the FED3 mode was changed to Timed Feed (FR and ABA mice only) and mice were fed from FED3 for 3 hours at the beginning of the dark cycle. Feeding data was collected on an SD card and further analyzed in R Studio.

### FED3 analysis

Pellet intake was measured across days of the ABA paradigm in FR and ABA mice using both cumulative and binned measurements. Binned measurements were determined for 15 minute intervals across the paradigm. Interpellet intervals were calculated in R Studio by subtracting the timepoints between individual pellet removals from the FED3 device. Pellets were classified as part of the same meal if they were retrieved within 130 seconds of another pellet (>1 pellet/meal requirement).

### Viral vectors

AAV1-hSyn-Flex-GCaMP6s (Addgene, 100845) was used for *in vivo* fiber photometry recordings of AgRP-expressing ARC neurons. AAV8-hSyn-Flex-mCherry (Addgene, 50459) and AAV8-hSyn-Flex-hM3Dq-mCherry (provided by NIEHS) were used for experiments designed for chemogenetic activation of AgRP neurons.

### Viral injections and optical fiber implants

Stereotaxic injections were performed as previously described^24^. Briefly, mice were anesthetized with isoflurane and placed in a stereotaxic frame (Stoelting’s Just for Mouse) and provided with analgesia (meloxicam, 0.5 mg/kg). Following a small incision on top of the skull and skull leveling, a small hole was drilled for injection. A pulled-glass pipette (20-40mm tip diameter) was inserted into the brain at coordinates aimed at the ARC (AP: −1.40, ML: +− .25), and 200 nL of virus was injected at two depths (DV: −5.75 and −5.65) using a micromanipulator (Grass Technologies, Model S48 Stimulator, 25 nL/min). For *in vivo* photometry experiments, an optic-fiber cannula (core = 400 um; 0.48 NA; M3 thread titanium receptacle; Doric Lenses) was implanted directly over the ARC (AP: −1.40, ML: +0.25, DV: −5.55) following virus injection. Fibers were fixed to the skull using C&B-Metabond Quick Adhesive Cement and dental acrylic. Following surgery, mice were placed on a heating pad and allowed to recover before single housing mice until further experimentation. All experiments were performed at least 2 weeks after stereotaxic surgery to allow for adequate viral expression and surgery recovery.

### In vivo fiber photometry set-up

To excite GCaMP6s, <20 uW blue LED light at 470 nm was driven by a multichannel hub (Thorlabs), modulated at 211 Hz, and subsequently delivered to a dichroic mini cube (FMC5, Doric Lenses) that was connected with optic fibers and a rotary joint (FRJ 1×1, Doric Lenses) to the optic cranial implant of the mouse. GCaMP6s calcium GFP signals were collected through the same fibers and dichroic minicube into a Femtowatt Silicon Photoreceiver (2151, Newport). Digital signals were subsequently demodulated, amplified and collected through a lock-in amplifier (RZ5P, Tucker-Davis Technologies (TDT)). LED modulation and data collection were performed using Synapse (TDT), exported via OpenBrowser (TDT) and analyzed in R Studio.

Freely-moving in vivo fiber photometry screening. To test for similar AgRP^GCaMP6s^ expression in all mice, mice were screened for viral and optic fiber hits. Mice were habituated to handling and hookup to the fiber patch cord for at least two days before screening. 18 hours prior to screening, food was removed from cages and mice were fasted overnight (during the dark cycle). On the morning of the screening day, mice were hooked up to the fiber patch cord, placed back in their homecage and were allowed to habituate to the set-up for at least three minutes, at which point recordings began. After at least two minutes of baseline recordings in the fasted state, a pellet of standard chow diet was placed on the cage bottom. Recordings continued for at least two minutes following pellet drop. Synchronized high-definition videos were recorded for time-locked data analysis in Synapse. For analysis, data was downsampled to 8 Hz and signals were normalized by z-score to the first minute of the recording (−120s → −60s) using R Studio.

Freely-moving in vivo fiber photometry during ABA. Mice were allowed to recover from the initial screening experiments for at least one week. Mice were then placed in one of three experimental cohorts (as described above) and housed in PhenoTyper cages (Noldus) containing a voluntary running wheel (locked, FR mice; unlocked, ABA and Activity mice), with the FED3 and standard chow on the ground. Fiber patch cord habituation was carried out in the light cycle for at least three days prior to Day 0 of the paradigm, during which mice were hooked up to the patchcord for at least 20 minutes. On both Day 1 and Day 4 of the paradigm, mice were hooked up to the fiber patch cord during the light cycle and were allowed to habituate for at least one hour before recordings began. Recordings began at least 15 minutes before the beginning of the dark cycle, allowing for detection of the first bout of feeding in FR and ABA mice. Food pellet dispense and retrieval were determined by TTL pulses driven by FED3 that were detected and time-locked to photometry recordings using Synapse. Recordings continued for one hour into the dark cycle, at which point recordings stopped and mice were unhooked from the fiber patch cord. Analysis was performed by downsampling recordings to 8 Hz and subsequently calculating perievent signals around pellet retrieval using NeuroExplorer (Plexon). Z-score calculations were performed with the mean and standard deviation of the first 5 seconds (−10 → −5) around each trial using R Studio.

### AgRP 3Dq screening

Following surgery recovery, AgRP^mCh^ and AgRP^3Dq^ mice were habituated to i.p. injections (100 uL saline) for at least three days. After habituation, all food was removed from the homecage of mice at the beginning of the light cycle, and one pre-measured chow pellet was provided to mice. At the same time, mice were injected with either vehicle (1U/g BW, 10% β-cyclodextrin) or clozapine n-oxide (CNO; 1 mg/kg in 10% β-cyclodextrin) in a crossover design with at least three days in between crossover. Food intake was measured at one hour after injection.

### AgRP activation experiments - Days 1-7

AgRP^mCh^ and AgRP^3Dq^ mice were allowed to recover for at least one week following screening experiments, and then were separated into one of three groups (Activity, FR, ABA) at least three days prior to the beginning of the behavioral paradigm. Mice were habituated to i.p. injections (100 uL saline) at least three days prior to Day 0 of the paradigm. Beginning on Day 0, all mice were injected with vehicle (Day 0; 1U/g BW, 10% β-cyclodextrin) or CNO (Days 1-7; 1 mg/kg) 15 minutes before the onset of the dark cycle. Behavioral paradigms were the same as described above in “Behavioral Paradigm.”

### AgRP activation experiments - Days 4-7

AgRP^mCh^ and AgRP^3Dq^ mice were all placed on the ABA behavioral paradigm, with food access provided by FED3 devices. Mice were habituated to i.p. injections (100 uL saline) at least three days prior to Day 0 of the paradigm. All mice were injected with either vehicle (Days 0, 1, 2, 3; 10% β-cyclodextrin) or CNO (Days 4, 5, 6, 7; 1 mg/kg in 10% β-cyclodextrin), and were exposed to the same behavioral paradigms as described above in “Behavioral Paradigm.”

### Perfusion and histology

Following experiment completion, mice were terminally anesthetized using chloral hydrate (Sigma-Aldrich) and transcardially perfused first with phosphate-buffered saline (PBS) followed by 10% neutral buffered formalin (Fisher Scientific). Brains were removed, post-fixed, and dehydrated in 30% sucrose before sectioning into 30-50 um slices using a freezing sliding microtome (Leica Biosystems). Coronal sections were collected in four series and stored at 4°C for immediate use, or −20C for long-term storage in antifreeze solution (1:4 glycerol, 1:4 ethylene glycol, 1:2 PBS). Slices were mounted with a mounting medium containing DAPI (Vectashield) and images were captured using a 10X objective on a Zeiss Observer Z1 confocal microscope.

### Statistical analysis

Paired t-tests, unpaired t-tests, one-way ANOVAs followed by Bonferroni post-hoc tests (if applicable), two-way ANOVAs followed by Bonferroni post-hoc tests (if applicable), or two-way mixed ANOVAs followed by Bonferroni post-hoc tests (if applicable) were calculated using R Studio as appropriate. Normality and homogeneity of variances were tested and, if necessary, accounted for using the Shapiro-Wilk and Levene’s tests, respectively. Significance was determined for p<0.05.

## Results

### Food Restriction with voluntary running wheel access exacerbates negative energy balance

Previous studies have demonstrated that body weight loss occurs in rodents in contexts of time-restricted food access in combination with access to a running wheel to varying degrees^19,20,45–47,19,20,45^. To recapitulate this behavioral paradigm, we placed wildtype (WT) mice in one of three cohorts: Activity, Food Restricted (FR), or Activity-Based Anorexia (ABA). Activity mice were provided *ad libitum* food throughout the experiment (thus serving as a control for voluntary wheel running activity, Fig. 1a). A separate control cohort consisted of FR mice that were food restricted and provided with a locked running wheel in their homecage (Fig. 1b). To measure the effects of both voluntary wheel running and food restriction, ABA mice were provided with a running wheel and were food restricted at the same times as FR mice, beginning on Day 0 of the paradigm (Fig. 1c).

**Figure 1.**
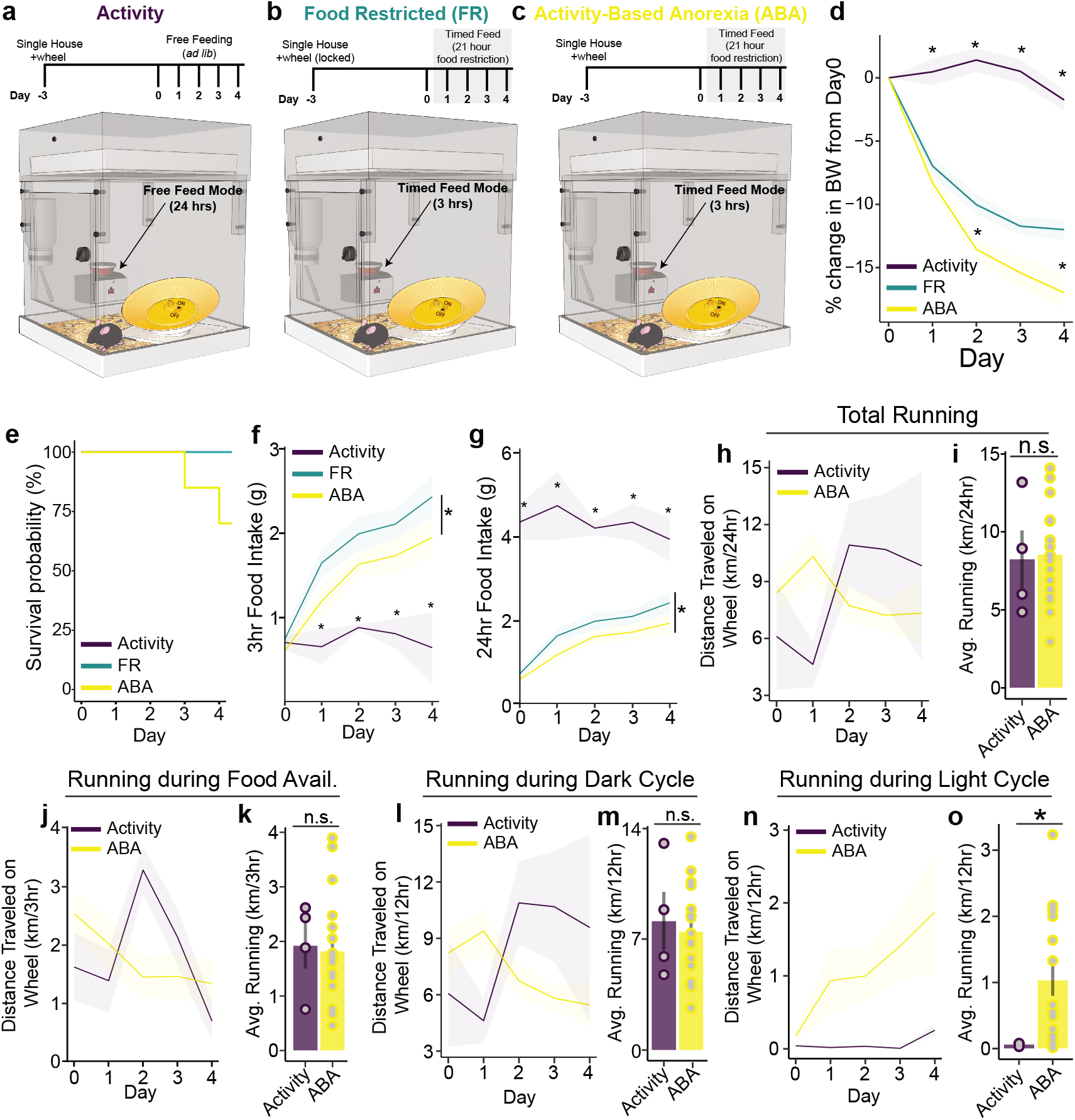
Food restriction with voluntary running wheel access causes activity-based anorexia. (a-c) Schematic of experimental set-up, in which mice are trained on a wireless voluntary running wheel in their homecage (Activity (a), Activity-Based Anorexia, ABA (c)) or a locked running wheel (Food Restricted, FR, b). Following wheel habituation, ABA and FR mice begin a 21-hour food restriction paradigm, in which food is available the first three hours of the dark cycle (beginning after the first three hours of the dark cycle on Day 0). (d) ABA mice lose more body weight than FR mice (Day 2 p=0.019, Day 4 p=0.046), whereas body weight of Activity mice remains constant. (e) Survival curves demonstrate that unlike Activity and FR mice, a portion of ABA mice need to be removed from the paradigm due to excessive body weight loss (>23%). Both ABA and FR mice eat more than Activity mice in the 3-hour period of food availability over the course of the paradigm (p values from Days 1-4 in order: Activity vs. FR p=0.023, p=0.006, p=0.015, p=0.007; Activity vs. ABA p=0.344 (n.s.), p=0.0647 (ns), p=0.103(ns), p=0.0324) (g). While ABA mice exhibit less food intake than FR mice during food availability (p=0.026), *ad libitum* fed Activity mice eat more than both groups over the entire 24 hour period (Activity vs. ABA/FR Days 0-3 p<0.0001; Activity vs. FR Day 4 p=0.026; Activity vs. ABA Day 4 p=0.002). ABA mice display no difference than *ad libitum* fed Activity controls in (h-i) total voluntary running, (j-k) running during the three hour window of food availability for ABA mice, or (l-m) running during the dark cycle but (n-o) exhibit increased wheel running during the light cycle (p=0.0009). Data for i, k, m, o averaged from days 0-4. *Wheel: n=4, FR: n=10, ABA: n=12-17*. Significance determined by p<0.05 using a two-way mixed ANOVA followed by Bonferroni post-hoc if appropriate (d, f-g, h, j, l, n) or an unpaired t-test (i, k, m, o). Data represented as mean ± SEM.

Similar to previous studies, mice in the Activity cohort maintain their body weight over the course of the paradigm, whereas FR causes mice to lose weight that stabilizes after Day 3 (Fig. 1d). In comparison, ABA mice exhibit weight loss that is more profound than FR mice (Fig. 1d). To this point, a proportion of ABA mice had to be removed from the behavioral paradigm due to significant body weight loss (>23%), whereas no mice in the Activity and FR cohorts were removed from study (Fig. 1e). Further analysis of energy balance parameters demonstrates that while FR and ABA mice both increase feeding during 3-hour food access over the course of the paradigm (two-way ANOVA, effect of time) in comparison to *ad libitum* fed Activity cohorts (Fig. 1f), ABA mice eat less than FR mice. Since Activity mice are provided *ad libitum* food, food intake in Activity mice is less than FR and ABA mice during the first three hours of the dark cycle, whereas 24-hr food intake is higher in Activity mice than either FR or ABA cohorts (Fig. 1g). This suggests that Activity mice maintain body weight by matching energy input with output, whereas ABA mice are unable to maintain body weight, at least in part due to diminished food intake relative to FR controls.

In concert with food intake analyses, we investigated the voluntary wheel running activity in both Activity and ABA mice. Although ABA mice are food restricted and losing body weight, their total wheel running activity is no different from *ad libitum* fed Activity controls (Fig. 1h-i). Similarly, despite food availability only during the first three hours of the dark cycle for ABA mice, running during this time period is similar to Activity controls (Fig. 1j-k). Since mice are typically more active during the dark cycle, we investigated whether Activity and ABA mice differed in their wheel running activity in relation to circadian cycles. The majority of time spent wheel running in Activity mice was during the dark cycle, and ABA mice ran a similar amount as Activity mice during this time period (Fig. 1l-m). Surprisingly, ABA mice ran significantly more than Activity mice during the light cycle, a period when Activity mice run on the wheel very little (Fig. 1n-o). This suggests that despite sustained body weight loss due to hypophagia, ABA mice voluntarily engage in wheel running activity at comparable (and sometimes elevated) levels as *ad libitum* fed mice.

### ABA mice have dysregulated feeding behavior

Previous studies interrogating the role of maladaptive feeding in ABA progression have been inconclusive, potentially due to different behavioral paradigms (e.g., methodology of food intake measurement, length and/or circadian timing of food availability)^34,41,45,48,49^. We suspected that this was also, at least in part, due to the fact that food intake is oftentimes more nuanced than the measurement of total intake. To address this limitation, we used a Feeding Experimentation Device (FED3) in a subset of FR and ABA WT mice, which was designed to accurately measure food intake in mice by dispensing 20 mg grain pellets and recording mouse retrieval over time (Fig. 2a-b)^44^. With this qualitative and quantitative approach, we demonstrate with more specificity the timepoints that ABA mice differ in their cumulative food intake in comparison to FR controls. 15-minute binned analyses across all days of the paradigm demonstrate that both FR and ABA mice ate most of the food at the beginning of food access (Fig. 2c, two-way ANOVA, main effect of time). However, ABA mice ate at a slower rate than FR controls, thus contributing to decreased total intake over time by Day 3 of the paradigm (Fig. 2d). We also identified that both groups of mice had additional smaller bouts of feeding at later time points during the 3-hour feeding period (e.g. timepoints 75-105 min in FR mice, Fig. 2c). Thus, we hypothesized that FR and ABA mice might differ in meal patterns.

**Figure 2.**
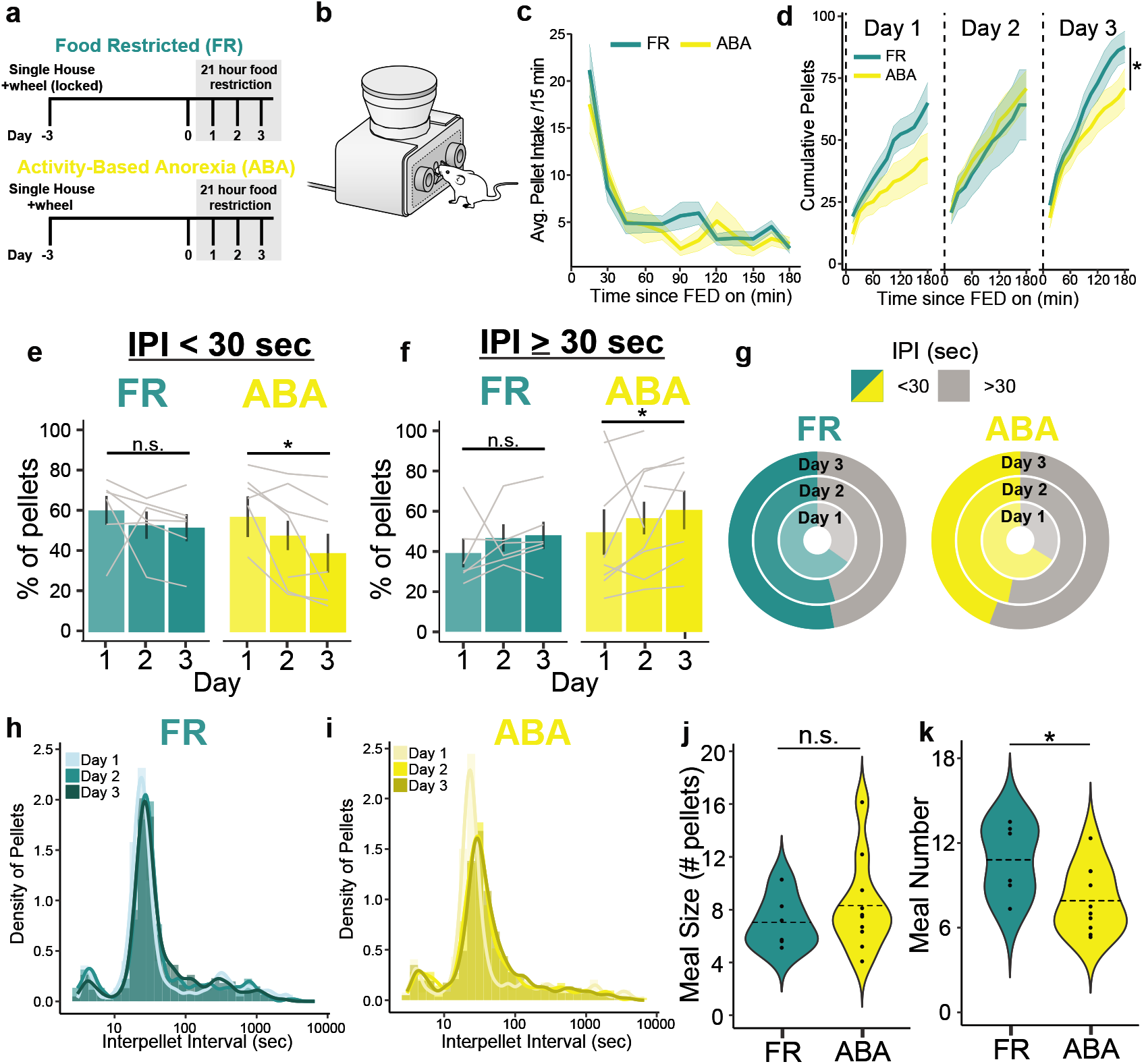
ABA mice demonstrate decreased food intake due to increased interpellet interval retrieval and diminished meal number. (a-b) Precise measurements of 3 hour feeding behavior was performed in FR and ABA mice using the Feeding Experimental Device (FED3, (b) throughout the paradigm. (c) 15-minute binned pellet intake illustrates feeding behavior over the course of the three-hour feeding window in FR and ABA mice. (d) Cumulative pellet intake over the course of feeding indicates ABA mice eat less than FR mice by Day 3 (p=0.026). e-g) The percentage of pellets eaten with an interpellet interval (IPI) <30 seconds (e) or IPI ≥ 30 seconds (f) is unaltered for FR mice across days of the paradigm, whereas ABA mice decrease and increase the frequency of pellets eaten with these IPIs, respectively (p=0.018). (g) Binned pellets from all mice within FR or ABA cohorts further exhibits progressively greater retrieval times in ABA mice (yellow/gray, right), compared to FR mice (green/gray, left). h-i) Kernel density estimation (KDE) curves indicate that FR mice maintain similar IPI frequencies throughout multiple days of food restriction, whereas the KDE curve of IPI in ABA mice shifts right after the first day of food deprivation. j-k) While meal size is comparable between FR and ABA mice (j), meal number is decreased in mice on the ABA paradigm compared to FR mice (k; p=0.049). Significance determined by p<0.05 using a two-way mixed ANOVA followed by Bonferroni post-hoc if appropriate (c,d), a one-way ANOVA followed by Bonferroni post-hoc if appropriate (e,f), or an unpaired t-test (j,k). *FR n=6; ABA n=7-10.* Data represented as mean ± SEM.

To accurately detect meal size and meal number, we first measured the interpellet interval (IPI) in both cohorts of mice, defined by the time (in seconds) in between each pellet retrieval from FED3 by the mouse. Previous reports suggest that the majority of pellets eaten from FED3 are consumed with an IPI <30 sec^44^. Thus, we measured the percentage of pellets eaten with an IPI <30 sec or ≥30 sec in FR and ABA mice across days of the paradigm (Fig. 2e-g). While FR mice do not alter the percentage of pellets eaten with an IPI <30 sec (Fig. 2e) or ≥30 sec (Fig. 2f) from Day 1 through Day 3 of the paradigm, ABA mice shift their food intake towards eating less percentage of pellets with an IPI <30 sec over the course of the paradigm (Fig. 2e), and more with an IPI ≥30 sec (Fig. 2f). Similarly, on Day 1 of food restriction, both FR and ABA mice show a kernel density estimation (KDE) peak in the relative IPI around 30 seconds (Fig. 2h-i, faint green/yellow). This KDE peak is maintained in the FR mice across Days 1-3, further suggesting that mice do not significantly alter how frequently they retrieve pellets over the course of multiple days of food restriction. In contrast, ABA mice shift the IPI KDE curves to the right on Day 2 and Day 3, suggesting more frequent increases in between each individual pellet retrieval as the behavioral paradigm progresses. These analyses demonstrate that ABA mice consume lower amounts of food than FR mice due to increased duration in between individual pellet retrieval.

### Meal number is decreased in ABA mice

Since IPI analyses (Fig. 2e-i) suggested that ABA mice alter IPI timing, we investigated whether this was reflected in either meal number or meal size in ABA mice compared to FR controls. While KDE peaks were observed with an IPI ~30 seconds on Day 1 across groups, we intended to determine meals based on the mean IPI of pellet consumption, which was ~IPI ≤130 seconds in FR mice (Fig. 2h-i), to most accurately detect meal bouts. Thus, pellets were classified as part of the same meal if they were within 130 seconds of one another. While ABA and FR mice demonstrated similar meal sizes (Fig. 2j), ABA mice had fewer meals than FR controls (Fig. 2i). This suggests that ABA mice have decreased feeding due to fewer initiation of meals rather than altering the amount of intake after a meal has started highlighting a behavioral distinction in ABA mice that potentially drives body weight loss.

### Feeding-induced AgRP inhibition is dysregulated in ABA mice

We next sought to identify whether hunger-promoting neurons were affected by food intake during the ABA paradigm. In particular, agouti-related peptide (AgRP)-producing neurons in the arcuate nucleus (ARC) have been well characterized for their ability to sense energy status via peripheral hormones and subsequently promote food intake across satiety conditions^24–26,50^. To this point, AgRP neurons are active in fasted mice, and are robustly inhibited upon food detection and durably suppressed contingent on subsequent caloric consumption^30,32,51^. Furthermore, abnormalities in AgRP mRNA and peptide levels have been described in ABA conditions, suggesting potential abnormalities at the level of AgRP neurons in this paradigm^40,41^. While previous studies demonstrated that AgRP neurons are modestly inhibited following exercise in food deprived states, no study to date has investigated the activity of this population during feeding in ABA. Given that AgRP neurons are well described to control food intake (more so than exercise) we assessed AgRP activity during feeding across ABA progression.

To measure the response of AgRP neurons during food access across behavioral conditions, we used *in vivo* fiber photometry in freely behaving *AgRP-iCre* mice (Fig. 3a-b). Before assigning mice to specific groups, we performed a baseline screening experiment aimed at testing injection and fiber placements during a fast-refeed (Fig. 3c). In line with previous literature, AgRP neurons were rapidly inhibited in response to food presentation in the fasted state^30,32,52^, and this response was similar across all mice that were subsequently divided into one of three behavioral groups (Activity, FR, ABA; Fig. 3c). This suggests that any differences observed in response to food intake during the behavioral paradigms are not due to targeting differences.

**Figure 3.**
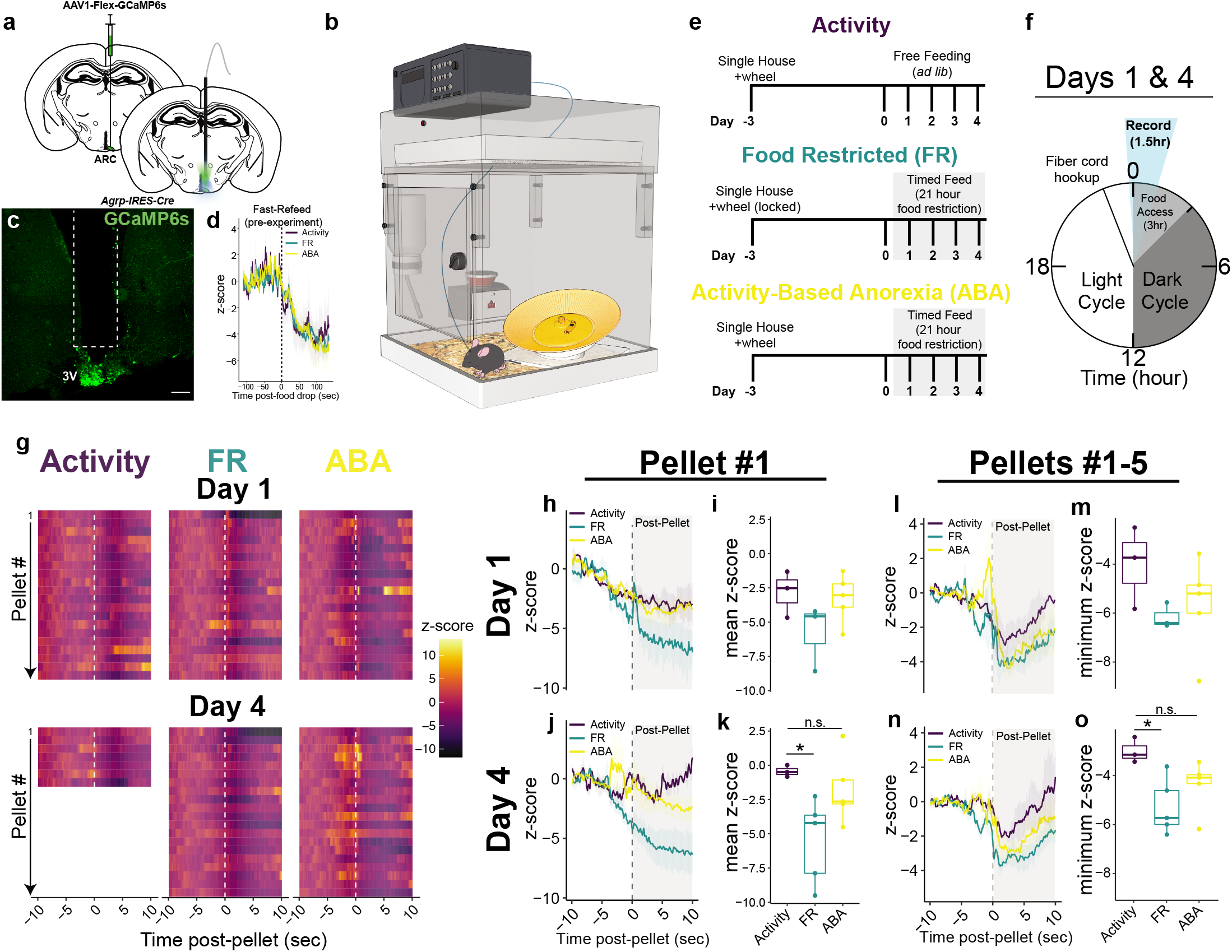
AgRP neuronal inhibition during food intake is dysregulated in ABA mice. (a-b) *Agrp-ires-Cre* mice were injected with AAV1-hSyn-Flex-GCaMP6s and a fiber placed above the ARC to record AgRP calcium signaling in freely-moving photometry experiments. (c) AgRP neuronal inhibition is similar across all behavioral groups in response to food presentation (t=0) in the fasted condition before being placed in experimental cohorts. (d) Behavioral set-up, in which mice are housed in phenotyper cages containing FEDs and voluntary running wheels, with AgRP calcium activity monitoring via fiber photometry patch cords that allow for free movement. (e,f) Mice in Activity, FR, or ABA conditions were hooked up to the fiber patchcord at least 1 hour before experiments began, and recordings began prior to food availability (f, blue). (g) Example raster plots of a mouse’s normalized AgRP calcium-induced activity across all 3 behavioral paradigms aligned to individual pellets eaten on Day 1 (top) or Day 4 (bottom). (h-o) Retrieval of either the first pellet (h-k) or the first five pellets (l-o) significantly decreases AgRP population activity in FR mice compared to *ad libitum* fed Activity controls (pellet 1 Day 4 Activity vs. FR p=0.0479; pellets 1-5 p=0.0278), whereas this decrease is not observed in ABA mice (k; averaged across t=0-10sec of Post-Pellet, m,o: minimum z-score across t=0-10sec of Post-Pellet). Significance was determined by p<0.05 using a one-way ANOVA followed by Tukey post-hoc if necessary (i, k, m, o). *Activity n=3*, *FR n=3-4; ABA n=5*. Data represented as mean ± SEM (h, j, l, n) or as a boxplot with line at median ± minimum/maximum (with outliers) (i, k, m, o); scale bar = 200 μm.

Previous studies utilizing fiber photometry approaches in the AgRP population have primarily focused on the effect of food presentation on AgRP activity over the course of 5-45 minutes, and thus potentially missed more nuanced alterations in AgRP activity. To this point, AgRP neurons can be modulated on shorter timescales, such as when aligned to individual bouts of food presentation and/or intake^31,52^. Thus, we sought to measure the effect of individual pellet intake on AgRP activity across intake using the FED3 devices. To probe the effects of food restriction on AgRP activity during ABA, photometry recordings were performed in homecages throughout the duration of the experiment (Day −3→Day 4, Fig. 3d) limiting the effects of stress on the animals. Behavioral paradigms were carried out as described in WT mice (Fig. 1), in which Activity mice had *ad libitum* access to food (provided from FED3 devices), and FR and ABA cohorts had access to food for three hours at the onset of the dark cycle beginning on Day 0 (Fig. 3e). Photometry recordings were performed before the start of the dark cycle (when food was available for FR and ABA mice) on Day 1 and Day 4 (Fig. 3f). Using this approach, we were able to perform perievent analyses on AgRP neuronal activity aligned to individual pellet intake (Fig. 3g). This analysis demonstrates that food intake (Pellet #1) significantly decreased AgRP activity in FR mice in comparison to *ad libitum* fed Activity controls by Day 4 of the paradigm; this inhibition was absent in ABA mice (Fig. 3h-k). Similarly, averaging AgRP neuronal activity dynamics across intake of the first five pellets demonstrates that AgRP neuronal inhibition in ABA mice more closely mimics that observed in Activity mice by Day 4 of the paradigm (Fig. 3l-o). This suggests that food intake-induced AgRP inhibition is disrupted during ABA, and that AgRP activity in ABA mice is more similar to that of *ad libitum* fed Activity mice than animals in a negative energy state (FR).

### Chemogenetic AgRP activation ameliorates body weight loss in ABA mice

Artificial AgRP activation, using either chemogenetic or optogenetic approaches, has repeatedly been demonstrated to increase food intake in rodents across multiple behavioral contexts^24–28,53–55^. Feeding-induced AgRP neuronal inhibition was dysregulated in ABA mice, suggesting the possibility that food intake in ABA mice does not relieve the negative valence associated with AgRP neuronal activation. We posited that this continued aversion prevents future food intake. Thus, we next sought to determine if driving AgRP neuronal activity could alleviate ABA development and/or reverse ABA progression by driving food seeking and food intake. While a recent study demonstrated that activation of AgRP neurons during exercise (without food present) does not facilitate future food intake (and actually promotes hyperactivity), we intended to maximize translational relevance and recapitulate decision-making in AN patients. Thus, we activated AgRP neurons during a period of behavioral choice (i.e. food intake vs. voluntary activity). To achieve this, we virally expressed hM3Dq (3Dq), a designer receptor exclusively activated by designer drugs (DREADDs) or a control fluorophore (mCherry, mCh) in AgRP neurons using *AgRP-iCre* mice (Fig. 4a-b). Initial screening experiments designed to test viral hit sites demonstrate that all cohorts of 3Dq-injected mice (Activity-3Dq, FR-3Dq, ABA-3Dq) similarly increase light-cycle feeding upon i.p. injection of CNO (the artificial ligand for the 3Dq receptor) when normalized to i.p. vehicle injections (Fig. 4c, “3Dq”). In contrast, control mice (Activity-mCh, FR-mCh, ABA-mCh) do not increase feeding in response to CNO injection (Fig. 4c, “mCh”). This suggests that all cohorts of experimental 3Dq mice are capable of increasing feeding to similar degrees in response to CNO administration.

**Figure 4.**
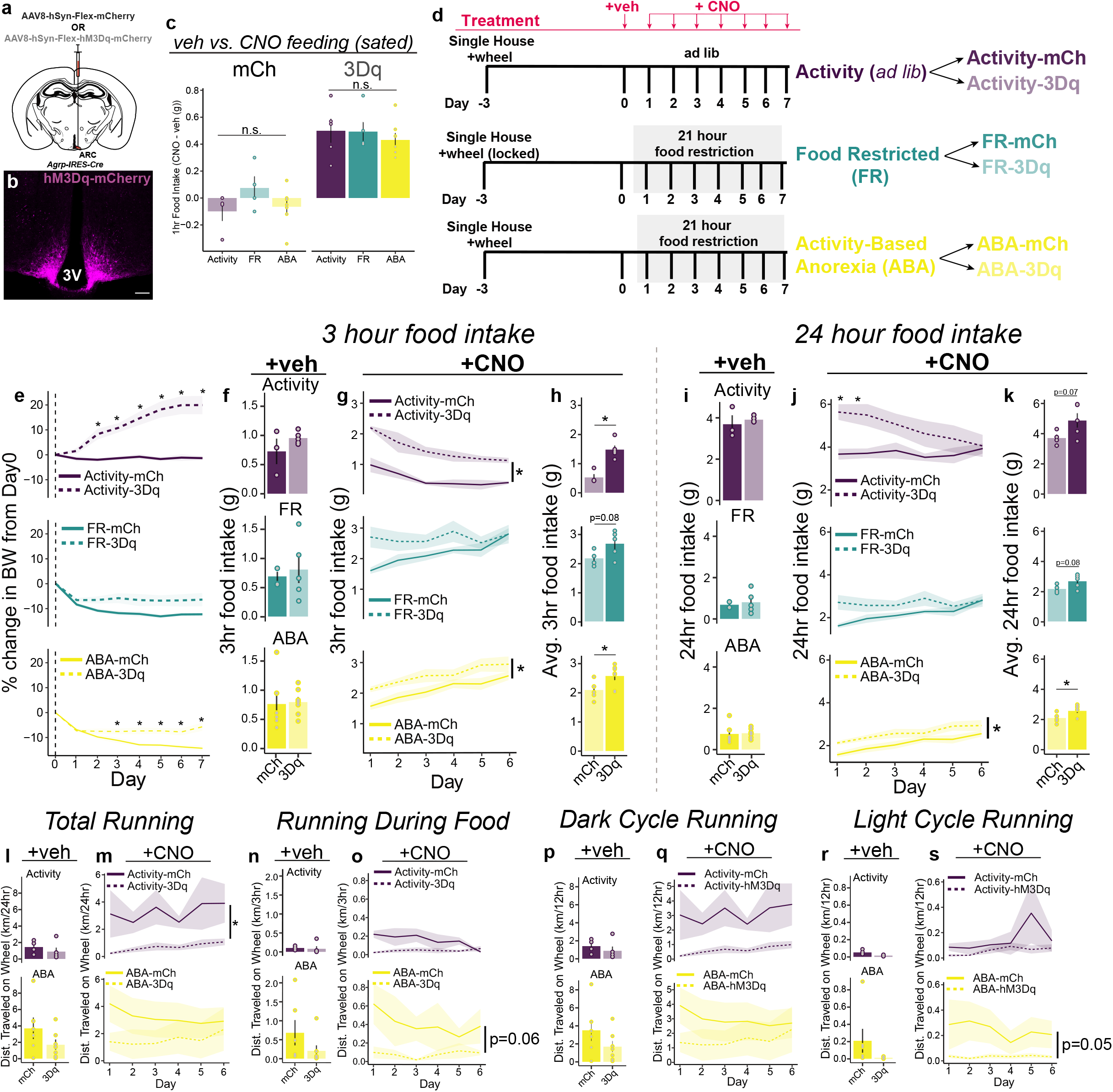
AgRP activation ameliorates ABA development. (a) AgRP neurons were artificially activated in *AgRP-iCre* mice via expression of 3Dq (using injection of AAV8-hSyn-Flex-hM3Dq-mCherry; 3Dq) and compared to control mice injected with AAV8-hSyn-Flex-mCherry (mCh). (b) Representative image of mCherry expression in the ARC of *AgRP-iCre* mice injected with AAV8-hSyn-Flex-hM3Dq-mCherry. (c) Initial viral hit site screening experiments across all mice before they were separated into experimental cohorts indicates that CNO induces food intake (normalized to vehicle injections) similarly across groups in 3Dq-expressing mice, with minimal food intake observed in mCh mice. (d) Outline of experiment, in which three behavioral groups each had control (Activity-mCh, FR-mCh, ABA-mCh) and experimental groups (Activity-3Dq, FR-3Dq, ABA-3Dq). All mice were injected with vehicle on Day 0 of the paradigm. Following 3 hours of feeding at the onset of the dark cycle on Day 0, FR and ABA groups were food restricted, whereas Activity mice were continued on *ad libitum* feeding. On Days 1-7, all mice received injections of CNO (i.p., 1.0mg/kg) 15 minutes prior to the onset of the dark cycle. (e) 3Dq-mediated AgRP activation (dashed lines) increases BW compared to mCh controls (solid lines) in Activity (top, purple; p=0.001) and ABA (bottom, yellow; p=0.001) mice. (f) 3-hour and (i) 24-hour food intake is comparable between mCh and 3Dq cohorts in response to vehicle injections on Day 0. 3Dq-mediated AgRP activation increases (g-h) 3-hour and (j,k) 24-hour feeding in Activity and ABA mice compared to mCh controls (Activity (g): p=0.001, Activity (h): p=0.001, Activity (j): p=0.003; ABA (g & j): p=0.03, ABA (h & k): p=0.0278). (l) Total, (n) 3-hour, (p) dark cycle and (r) light cycle voluntary running activity is comparable between mCh and 3Dq cohorts in response to vehicle injections on Day 0. 3Dq-mediated AgRP activation in Activity mice decreases total voluntary running (m; p=0.013), with no significant effect on running during food availability (o), dark cycle running (q), or light cycle running (s). 3Dq-mediated AgRP activation in ABA-3Dq mice does not significantly alter running behavior across analyses (m, o, q, s), but does show a trend towards decreased running during food availability (o, p=0.06) and decreased light cycle running (s, p=0.05). Significance determined by p<0.05 using a one-way ANOVA (c), a two-way ANOVA followed by Bonferroni post-hoc if appropriate (e, g, j) a two-way mixed ANOVA followed by Bonferroni post-hoc if appropriate (m, o, q, s; Activity only), or an unpaired t-test (f, h, i, k, l, n, p, r). *Activity-mCh n=3-4; Activity-3Dq n=5; FR-mCh n=4; FR-3Dq n=5; ABA-mCh n=6; ABA-3Dq n=7.* Data represented as mean ± SEM

To test the effect of chemogenetic activation of AgRP neurons in ABA progression, each cohort of mice had both a control (mCh) and experimental sub-cohort (3Dq). All mice were administered vehicle on Day 0 of the paradigm 15 minutes prior to the onset of the dark cycle, whereas CNO was administered on Days 1-7 in all cohorts (Fig. 4d). Using this approach, we were able to decipher the effect of artificial AgRP stimulation in *ad libitum* fed Activity mice (Activity-mCh vs. Activity-3Dq), FR mice (FR-mCh vs. FR-3Dq), and ABA mice (ABA-mCh, ABA-3Dq). Indeed, activation of AgRP neurons increased body weight in *ad libitum* fed Activity mice (Fig. 4e, purple) and ABA mice (Fig. 4e, yellow). AgRP stimulation did not have a significant effect on the body weight of FR mice, likely due to ceiling effects of food intake during the three hour feeding window (Fig. 4e, green).

Since we initially determined that body weight loss is at least partly due to a reduction in food intake in ABA mice, we investigated if food intake was altered in AgRP-3Dq mice compared to their AgRP-mCh controls. While no change was observed in food intake during the first three hours of the dark cycle on Day 0 (pre-AgRP activation, Fig. 4f), significant increases in food intake were observed in both Activity-3Dq and ABA-3Dq mice in comparison to their mCh control cohorts when CNO was administered (Fig. 4g-h). Similarly, 24-hour intake was increased in Activity-3Dq mice in comparison to Activity-mCh controls, and this increase was particularly prevalent on the first two days of CNO administration (Fig. 4j). FR-3Dq mice did not demonstrate increased food intake in response to CNO administration compared to FR-mCh controls, suggesting that AgRP activation in food deprived conditions (absent access to a voluntary running wheel) is unable to increase feeding beyond the underlying homeostatic drive to eat in times of energy deficit. These results suggest that ABA body weight reversal as a result of AgRP activation is due, at least in part, to increased food intake.

Our initial behavioral tests indicated that ABA mice do not significantly alter total running in comparison to *ad libitum* fed Activity control mice, suggesting that voluntary wheel running activity in contexts of food restriction is likely a contributing factor to the body weight loss observed in ABA mice. Thus, we next investigated how chemogenetic AgRP activation alters voluntary wheel running activity in *ad libitum* fed and food deprived conditions in Activity and ABA mice, respectively. Voluntary running activity in Activity-3Dq and ABA-3Dq mice was unaltered in response to vehicle injections on Day 0 across all measured conditions (Fig. 4l, n, p,r) in comparison to Activity-mCh and ABA-mCh controls, respectively. While total voluntary wheel activity was decreased in Activity-3Dq mice compared to Activity-mCh mice, ABA-3Dq mice did not significantly decrease voluntary wheel activity (Fig. 4m, o, q, s). Taken together, these results suggest that AgRP activation in the context of *ad libitum* food with running wheel access increases body weight due to increased food intake and decreased wheel running. Similarly, AgRP activation in the context of food restriction and running wheel access decreases body weight loss primarily due to increased food intake.

### Chemogenetic AgRP neuronal activation nullifies ABA progression

Although our initial chemogenetic experiments were aimed to test the preventative capabilities of AgRP activation on ABA development, this approach has limited translational potential since most individuals exhibit ABA before intervention is possible. To test the capability of AgRP neuronal activation to rectify ABA, we performed AgRP activation after ABA progression had begun and mice had already lost significant body weight. Since we found that AgRP neuronal responses to food retrieval in ABA mice are more similar to that observed in *ad libitum* fed Activity mice than FR mice on Day 4 of the behavioral paradigm (Fig. 3j-k, 3n-o), we highlighted this time point as a potential intervention day. To test this, ABA-mCh and ABA-3Dq mice were initially injected with vehicle at the beginning of the ABA paradigm (Fig. 5a-b). Beginning on Day 4 of the paradigm, all mice received CNO injections, thereby activating AgRP neurons in ABA-3Dq mice following initial body weight loss (Fig. 5b). While ABA-mCh and ABA-3Dq mice did not differ in their initial body weight loss during vehicle injections, CNO administration in ABA-3Dq mice beginning on Day 4 was capable of rapidly and robustly improving body weight loss compared to ABA-mCh controls (Fig. 5c). This amelioration was partially due to increased food intake in ABA-3Dq mice compared to ABA-mCh controls during CNO administration days (Fig. 5d, Day 4-7), whereas food intake in response to vehicle injection was not different between groups (Fig. 5d, Days 0-3).

**Figure 5.**
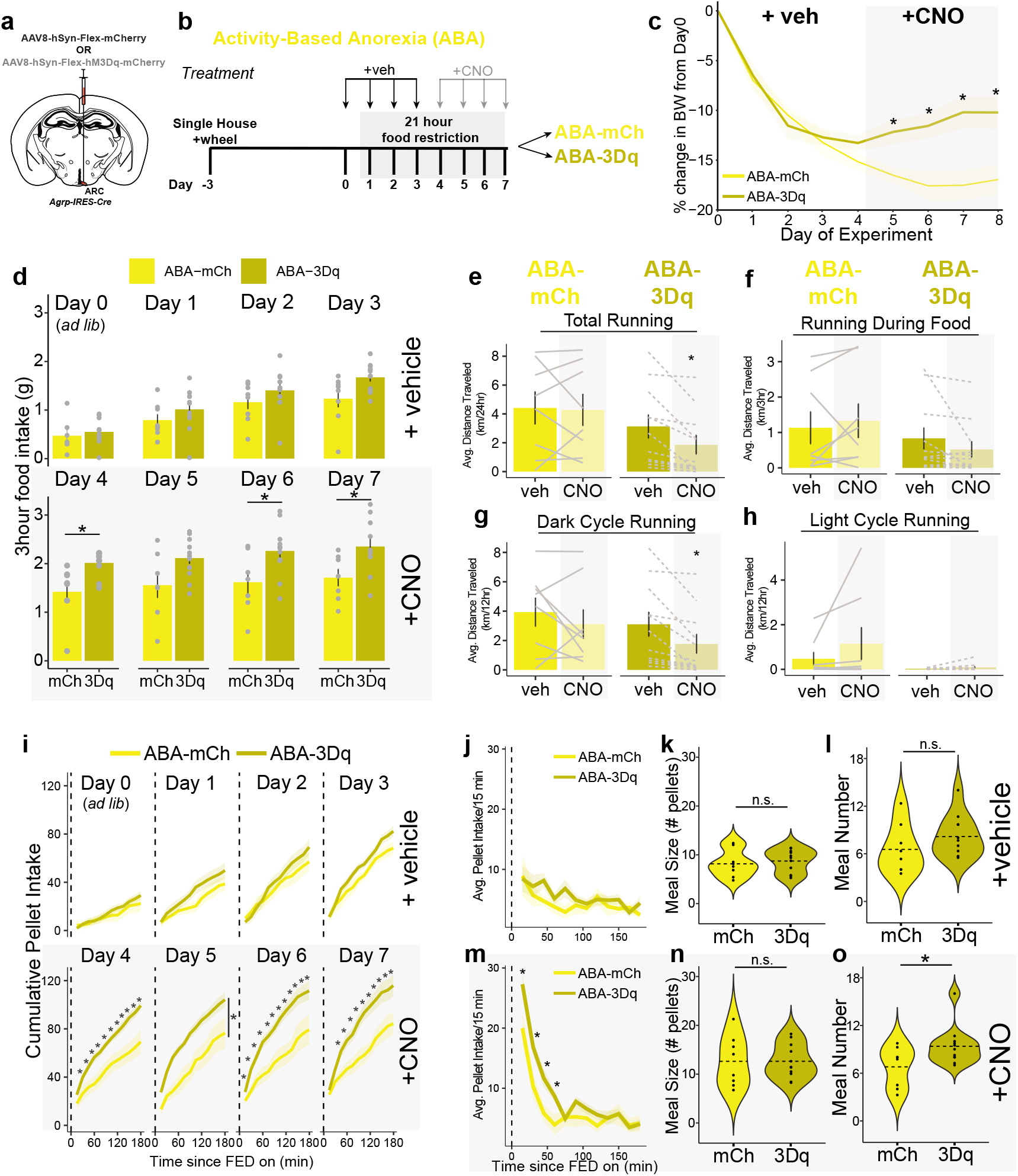
ABA progression is rectified by AgRP activation after initial body weight loss. (a) AgRP neurons were artificially activated in *AgRP-iCre* mice via expression of 3Dq (using injection of AAV8-hSyn-Flex-hM3Dq-mCherry; 3Dq) and compared to control mice injected with AAV8-hSyn-Flex-mCherry (mCh). (b) ABA paradigm where mice were injected with vehicle on Days 0-3 and CNO on Days 4-7 in all mice (ABA-mCh; ABA-hM3Dq). (c) While body weight change in ABA-mCh and ABA-3Dq mice is comparable on days following vehicle injection (Days 1-4; white background), ABA-3Dq, but not ABA-mCh, mice regain body weight the day following CNO administration (Days 5-8; grey background; Days 5-8 p=0.0145, p=0.0075, p=0.0064, p=0.0123). (d) Vehicle injections do not alter food intake between ABA-mCh and ABA-3Dq mice on Days 0-3, whereas CNO administration increases total feeding in ABA-3Dq mice compared to ABA-mCh controls (Day 4 p=0.0194, Day 6 p=0.0453, Day 7 p=0.0263). (e-h) CNO injections in ABA-mCh mice does not alter total running (e), running during food (f), dark cycle running (g), or light cycle running (h), while CNO-mediated AgRP activation in ABA-3Dq mice decreases (e) total (p=0.0075) and (g) dark cycle running (p=0.0073), but not (f) running during food or (h) light cycle wheel running (Days 4-7) compared to vehicle injected days. (i) Cumulative pellet intake and (j, m) 15-minute binned pellet intake was comparable in response to vehicle injections on Days 0-3 in ABA-mCh and ABA-3Dq mice (top). CNO injection increased both cumulative pellet intake (i, bottom; Day 4 (group x time): p=0.001, Day 5 (group): p=0.006, Day 6 (group x time): p= 0.026, Day 7 (group x time): p=0.007) and binned pellet intake especially during the first hour of food availability (m; p=0.015) in ABA-3Dq mice compared to ABA-mCh control mice. (k, n) Meal size was unaffected by either (k) vehicle or (n) CNO injections across groups. (l) Meal number was comparable in response to vehicle injections on Days 0-3. (o) CNO injection increased meal number in ABA-3Dq mice compared to ABA-mCh control mice (p=0.0413). Significance determined by p<0.05 using a two-way mixed ANOVA followed by Bonferroni post-hoc if appropriate (c) a two-way ANOVA followed by Bonferroni post-hoc if applicable (j, m, i), an unpaired t-test (d, k, l, n, o), or a paired t-test (e-h). *ABA-mCh n=7-8, ABA-3Dq n=10-11.* Data represented as mean ± SEM.

Concomitant with food intake measurements, we investigated whether AgRP activation altered running wheel activity after initial ABA progression. Indeed, total and dark cycle wheel running was decreased on days in which CNO was administered in ABA-3Dq mice compared to vehicle-injected days (Fig. 5e, G, ABA-3Dq). In contrast, all measurements of wheel running activity were unchanged between CNO and vehicle injection days in ABA-mCh mice (Fig. 5e-h, ABA-mCh). This suggests that body weight reversal in ABA-3Dq mice is partially due to both decreased wheel running activity and increased food consumption.

### Chemogenetic AgRP neuronal activation modifies meal number deficiencies in ABA mice

Our initial experiments uncovered that ABA mice initiate fewer meals than FR controls, thus highlighting a potential behavioral intervention method of increasing food intake in ABA conditions. To assess if AgRP activation-induced ABA amelioration was in part due to altered meal patterns, we further analyzed feeding data from FED3 devices used by ABA-mCh and ABA-3Dq mice in response to vehicle (Days 0-3) or CNO injections (Days 4-7). With this approach, we demonstrate that cumulative pellet intake was immediately increased upon the initiation of feeding in ABA-3Dq mice when CNO was administered on Days 4-7 of the paradigm, whereas this was not observed in response to vehicle injections on Days 0-3 (Fig. 5i). Binned pellet analyses further illustrate these findings, in which intake is significantly increased at t=15, 30, 45, and 60 minutes following food availability in CNO-injected ABA-3Dq mice compared to ABA-mCh controls (Fig. 5m). While meal size was unaltered in response to AgRP activation (Fig. 5n), CNO administration in ABA-3Dq increased meal number compared to ABA-mCh mice (Fig. 5o). These findings demonstrate that artificial AgRP activation reverses the meal number deficiency characterized by ABA, thus contributing to a rescue of ABA progression.

## Discussion

Studies have emphasized the behavioral complexities of ABA, and more recent reports have begun to unravel the neurobiology potentially underlying ABA^34,42,45–47,56,57^. Despite these advances, no study to date has identified a neural circuit capable of combating ABA development and/or progression by addressing both anorexia and hyperactivity. Here, we highlight the ability of AgRP neurons to override these maladaptive behaviors and subsequently coordinate body weight maintenance and survival. We first identify the feeding microstructure in mice displaying ABA and highlight meal number as the main feeding deficiency in ABA mice. We also demonstrate that AgRP neuronal inhibition in response to food intake is impaired in ABA mice, such that AgRP dynamics more closely resemble *ad libitum* fed conditions than animals in caloric deficit. We subsequently addressed these abnormalities in ABA mice using chemogenetic activation of AgRP neurons throughout ABA, and were able to alleviate ABA body weight loss via increased feeding. Finally, we demonstrate that AgRP activation is capable of mitigating ABA following the initial progression of body weight loss via increased food intake and decreased wheel running. Importantly, the reversal of ABA by AgRP activation is performed during food availability, clarifying the capability of AgRP neurons to promote food intake when activation is performed during a more physiologically relevant time (i.e. when endogenous AgRP neuronal activity is elevated).

AgRP activation in contexts without food is aversive; this negative valence associated with heightened AgRP activity is relieved upon food intake in healthy individuals^26,31^. While fasting-induced AgRP neuronal inhibition has been well documented in response to less than one day of fasting, no studies have demonstrated the effect of multi-day food restriction on AgRP neuronal activity. Here, we demonstrate that AgRP neurons are appropriately inhibited during food intake following one or four days of food restriction. However, this inhibition is lost in contexts with voluntary wheel running, suggesting that feeding-induced AgRP neuronal inhibition more closely resembles a sated state during ABA. These results are similar to the decreased inhibition of AgRP neurons observed in response to presentation of normal chow following exposure to an obesogenic diet^58,59^. Yet, in diet-induced obesogenic states, AgRP stimulation-induced standard chow intake is significantly blunted, whereas activation of AgRP neurons in ABA conditions strongly promotes chow intake. Thus, the capability of AgRP activation to trigger food intake is uniquely tailored to non-positive energy balance states.

While AgRP neurons have been well described for their ability to both sense peripheral energy signals and subsequently drive feeding behavior, few studies have measured their capability to coordinate maladaptive anorexic behaviors^24–26,51,60^. AgRP neurons are necessary for feeding (and subsequent survival) in adult mice, and are able to drive feeding across physiologic and behavioral contexts, including circadian cycles, social interaction, threat detection, and pain^27,29,61–64^. Moreover, manipulations of the melanocortin-3 receptor in the ARC (which is present on almost all AgRP neurons) has recently been shown to contribute to anorexic behaviors in mice^65,66^. Here, we identify for the first time the time scale required for AgRP neurons to be capable of driving food intake during ABA, a condition of unique behavioral adaptations, including hyperactivity. Activation of AgRP neurons in this study was performed during the initiation of the dark cycle, when mice typically begin eating. In contrast, activation of AgRP neurons during the light cycle in ABA mice without food present promotes hyperactivity and continues promoting body weight loss, highlighting the necessity of food availability to AgRP-mediated ABA reversal^34^. Moreover, AgRP neurons are inhibited following exercise in food-restricted states, suggesting that artificial activation of this population during restriction might drive behaviors intended to promote inhibition and alleviate the aversive nature of AgRP activation^33,34,53^.

A diversity of literature describes overall food intake in ABA, and yet no overarching consensus has been made on the microstructure of food intake during this condition. Here, we used specialized devices^44^ to detect nuanced feeding bouts more accurately in mice, and thus are able to highlight decreased meal number as a component of the decreased overall feeding observed during ABA. The peripheral and neural mechanisms underlying differences in meal size and meal number are largely unknown. While it is well documented that AGRP is elevated during hyperactive anorexia in humans and rodents, it is unclear if this elevation drives the altered meal patterns we observed^38–42^. Despite this elevation, we demonstrate that feeding-induced inhibition of AgRP neural activity is dysregulated during ABA. It is conceivable that these alterations in AgRP activity in response to food intake drive the observed behavioral abnormalities (i.e. decreased meal number and/or hyperactivity), but this has not yet been tested.

Taken together, we have ascertained the unique capability and circadian timeframe necessary for AgRP neurons to ameliorate ABA despite disrupted neural activity responses to food. While these studies measure bulk activity responses to food intake, subsequent studies will be required to identify if AgRP neuronal activity dynamics are heterogenous using single-cell imaging techniques. Furthermore, future studies aimed at characterizing the molecular changes in AgRP neurons following ABA have the potential to elucidate the cellular mechanisms coordinating these changes, and thus greatly advance the collective understanding of feeding behavior during states of negative energy balance.

## Acknowledgements

We would like to thank Dr. Alexxai Kravitz for the development of and extensive help with FED3 devices. We thank Dr. Bryan Roth for the CNO used in chemogenetic experiments. We also thank N. Martin and B. Gloss of the NIEHS Viral Vector Core for producing the AAVs used in chemogenetic experiments. We thank all members of Dr. Krashes’ lab for technical support and guidance during this project. This research was supported by the Intramural Research Program of the National Institutes of Health, the National Institutes of Diabetes and Digestive and Kidney Diseases (DK075088 to M.J.K. and DK075087-06 to M.J.K) and the Nancy Nossal Fellowship (NIH-NIDDK; to A.K.S.).

## Contributions

A.K.S. and M.J.K. designed experiments. A.K.S. and S.C.D performed initial experiments in WT mice. A.K.S, S.C.D., A.M.S., A.S. constructed FED3 devices used throughout experiments. A.K.S. performed chemogenetic and photometry experiments. A.K.S. performed analysis on all experiments. A.K.S. and M.J.K. wrote the manuscript with input from S.C.D.

---

**Supplementary Video 1.** *Example video of food restricted (FR) and activity-based anorexia (ABA) mouse behavior during food access.* A mouse on the FR paradigm (left) eats from a Feeding Experimentation Device (FED3) when food is accessible during a photometry recording after 21 hours of food deprivation. A simultaneous recording of a mouse on the ABA paradigm (also following 21 hours of food deprivation) illustrates that the ABA mouse chooses to voluntarily run on the wireless running wheel rather than eat chow pellets from the FED3.

## Notes

### Competing Interest Statement

The authors have declared no competing interest.

